# Insights into the performance of CusF as a solubility tag for recombinant protein expression

**DOI:** 10.64898/2025.12.08.693106

**Authors:** Igor P. Oscorbin, Maria A. Smertina, Maria S. Kunova, Maxim L. Filipenko

## Abstract

The metal-binding periplasmic protein CusF has been proposed as a bifunctional tag enhancing solubility of recombinant proteins and enabling purification using Cu affinity chromatography. However, evidence for its performance remains limited to a few model proteins. Here, we evaluated CusF as a solubility tag for two heterologous proteins: a putative poly(A)-polymerase from *Enterococcus faecalis* (Efa PAP) and the red fluorescent protein mCherry. The proteins were fused to CusF, expressed in *E. coli* BL21 (DE3) pLysS and Rosetta 2 (DE3) strains, and assessed for solubility and IMAC binding. Native Efa PAP was completely insoluble under all tested conditions, and fusion to CusF did not improve its solubility. Similarly, CusF-mCherry accumulated predominantly in the insoluble fraction, with only traces detectable in soluble lysates. Soluble CusF-mCherry did not bind Cu^2+^-charged IMAC resin, while moderate binding to Ni^2+^-charged resin was attributable to the vector-encoded His-tag rather than CusF. These results indicate that CusF does not universally enhance protein solubility and may not always bind Cu-based IMAC resin. Our findings expand empirical knowledge on solubility tag performance and emphasize the necessity of testing multiple tags to identify optimal strategies for recombinant protein production.

## 1. Introduction

Expression of recombinant proteins in *E. coli* is one of the most prominent methods in modern biotechnology, widely used in both fundamental studies and practical applications. After expression, recombinant proteins are typically isolated by various approaches, most commonly by chromatography. Despite several decades of protocol optimization, recombinant protein expression is still complicated by issues such as low yield and low solubility. Insufficient protein production often results in its inadequate amount for downstream studies or applications. Low protein solubility frequently necessitates denaturation followed by refolding, which not only complicates purification and increases its cost but also can lead to a loss of specific activity. Among the solutions proposed to overcome these challenges, protein tags remain among the most effective. These tags are small- or medium-sized proteins or protein domains fused to the protein of interest, such as maltose-binding protein (MBP) [1], glutathione S-transferase (GST) [2], SUMO [3], thioredoxin [4], NusA [5] and many others. During purification, tags can be removed by proteolytic cleavage to obtain native recombinant proteins. Protein tags are multifunctional, not only directly increasing yield and solubility but also directing recombinant proteins to the periplasm or culture medium (e.g., pelB) [6], facilitating purification through affinity interactions (e.g., MBP, GST), or even enhancing the functionality of fused proteins, as demonstrated for Sso7d-fused DNA polymerases [7]. Nevertheless, identifying the optimal protein tag remains largely empirical and must be determined individually for each recombinant protein.

Among available protein tags, CusF was recently proposed as a bifunctional metal-binding protein that increases solubility of fused proteins and binds Cu-charged IMAC resin. CusF is a small periplasmic *E. coli* protein (12.3 kDa, 110 aa) that binds Ag^+^ and Cu^+^ ions (K_d_ 38.5 ± 6.0 nM and 495 ± 260 nM, respectively) and transfers them to the CusCBA efflux pump for export across the outer membrane [8,9]. Together with the other proteins of the CusCFBA system, CusF protects *E. coli* cells from Cu and Ag toxicity. CusF was reported to be an efficient solubility tag for GFP expression in *E. coli*, outperforming MBP and GST, and to bind Cu^2+^-charged IMAC resin [10]. An enhanced CusF mutant, CusF3H, was later developed to bind His-charged IMAC through the addition of three N-terminal His residues [11]. However, data on the applicability of CusF for recombinant protein expression and purification remain limited to only a few examples [11,12], leaving room for further investigation.

Polyadenylation is an RNA modification involved in RNA decay in prokaryotes. Poly(A) tails serve as recognition signals for nucleases—PNPase, RNase II, and RNase R—which initiate RNA degradation from these tails. Poly(A) polymerase (PAP) [8] is primarily responsible for polyadenylation in *E. coli* [13]. Only a few PAPs have been characterized from other bacteria, including *Geobacter sulfurreducens* [14], *Pseudomonas putida* [15], and *Bacillus subtilis* [16]. However, only the PAP from *G. sulfurreducens* has been cloned, while the other two were purified directly from native hosts. Thus, the diversity of bacterial PAPs remains greatly understudied, even as these enzymes gain increasing interest due to their use in manufacturing mRNA therapeutics. It should also be noted that *E. coli* PAP is prone to aggregation and instability in low-salt buffers, making it a difficult target for purification [13,17].

Fluorescent proteins are indispensable for numerous applications, including studies of in vivo protein localization and dynamics, cell imaging, and reporting cellular processes. A broad palette of fluorescent proteins has been developed with varying quantum yields and emission wavelengths. They also serve as convenient tools for optimizing protein purification protocols, allowing rapid and visually detectable readouts. One of often used fluorescent proteins, mCherry is a far-red monomeric fluorescent protein engineered from DsRed and widely used for deep-tissue imaging. mCherry is mostly soluble in and expressed in high amounts in *E. coli* cells, but has low homology with GFP, being a good alternative to the latter for monitoring the purification efficacy.[18]

Here, we applied the small metal-binding *E. coli* protein CusF as a solubility tag for purification of a putative novel poly(A) polymerase from *Enterococcus faecalis*, a commensal bacterium of the human gastrointestinal tract, as well as for purification of the fluorescent protein mCherry. After cloning the fusion constructs, we expressed them in two *E. coli* strains and assessed their solubility and ability to bind Cu- and Ni-charged IMAC resins.

## 2. Results

### 2.1. Expression of a putative Efa PAP and its fusion with CusF

To identify new bacterial PAPs, we aligned the amino acid sequence of *E. coli* PAP-1 using Protein BLAST against proteins from *Bacilli*. The search criteria included a length of 400–600 amino acids and the presence of the characteristic bacterial PAP signature motif: [LIV][LIV]G[R/K][R/K]Fx[LIV]h[HQL][LIV]. Candidate proteins containing additional domains were excluded to avoid the presence of unrelated enzymatic activities. The retrieved putative PAPs were ranked based on their similarity to *E. coli* PAP-1, and from 223 candidates, we selected a putative PAP from *Enterococcus faecalis* (GenBank: EOE33521.1), taking into account the ecological niche of the host. *Enterococcus faecali*s is a non-motile, facultatively anaerobic coccus that resides as a commensal organism in the human gastrointestinal tract and is an opportunistic pathogen causing urinary tract infections, endocarditis, and bacteremia. It grows optimally at 6.5% NaCl, 37 °C, and in the presence of 40% bile [19]. Thus, genomic DNA from this species is relatively easy to obtain. The putative *E. faecalis* PAP (Efa PAP) is a 48.5 kDa protein (424 amino acids) with a low isoelectric point (pI 5.29), a strongly negative charge at pH 7.0 (–13.1) and low sequence similarity to *E. coli* PAP-1 (35.41%). The coding sequence of Efa PAP was cloned into the pET23a vector; however, recombinant Efa PAP was insoluble after expression in *E. coli* BL21 (DE3) pLysS and Rosetta 2 (DE3) strains at both 25 °C and 37 °C (Figure 1a and 1b).

**Figure 1.**
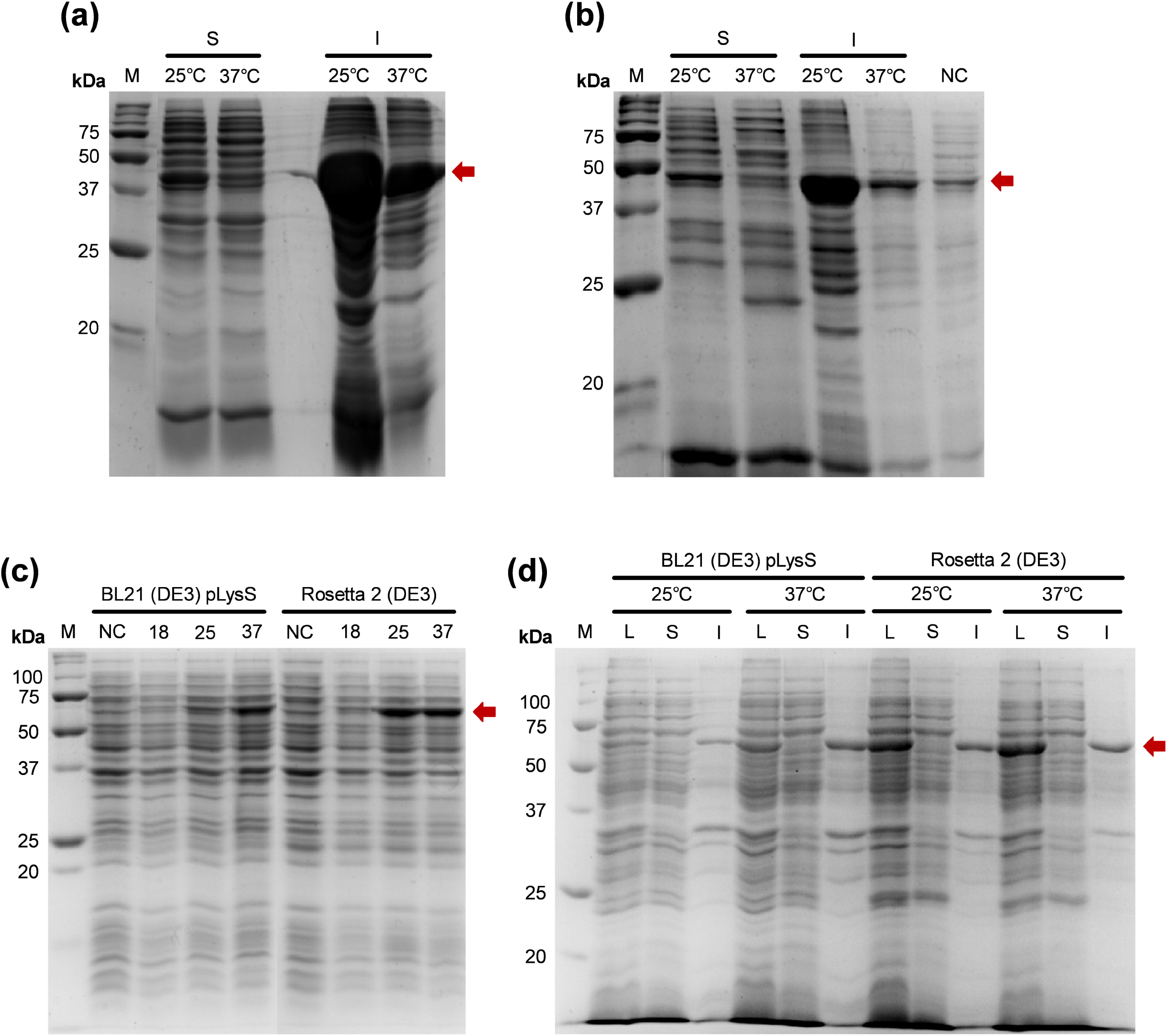
Expression of the putative *E. faecalis* PAP and its fusion with CusF. The proteins were expressed in *E. coli* strains BL21 (DE3) pLysS and Rosetta 2 (DE3) at various temperatures. M — Precision Plus Protein standards (Bio-Rad, Hercules, CA, USA). Red arrow marks the recombinant spectively. 25°C, 37°C —expression temperature, S — soluble protein fraction, I — insoluble proteins, NC — negative control before expression. (c) — Expression of the putative Efa PAP fused with CusF. 18, 25, 37 —expression temperature in °C, NC — negative control before expression. The strains are indicated above the respective lanes. (c) — Solubility of the putative Efa PAP fused with CusF. 25°C, 37°C —expression temperature, L — lysates after the expression, S — soluble protein fraction, I — insoluble proteins. The strains are indicated above the respective lanes.

To increase the solubility of Efa PAP, we fused the protein with CusF, a Cu-binding protein previously reported as a solubility tag capable of binding Cu-charged IMAC resin, thereby potentially improving both solubility and suitability for metal-chelate chromatography. The fusion construct was cloned into the pET36b vector and expressed in the same *E. coli* strains used for Efa PAP (Figure 1c and 1d). However, the CusF–Efa PAP fusion protein was also completely insoluble, regardless of the strain or induction temperature. It should be noted that we did not detect poly(A) polymerase activity under conditions similar to those used for *E. coli* PAP-1, either in lysates with Efa PAP or in lysates with the CusF–Efa PAP fusion protein (data not shown). Therefore, the possible impact of CusF on the specific activity of Efa PAP remains unknown.

### 2.2. Expression and binding with IMAC of a CusF-mCherry fusion

After the unsuccessful attempts to obtain soluble Efa PAP, we fused CusF to mCherry, a DsRed-derived fluorescent protein, as a model substrate similar to GFP—the protein originally used to demonstrate the applicability of CusF as a solubility tag. The CusF–mCherry fusion protein was expressed in *E. coli* BL21 (DE3) pLysS and Rosetta 2 (DE3) strains, and the resulting lysates were incubated with either Cu^2+^-charged or Ni^2+^-charged IMAC resin (Figure 2).

**Figure 2.**
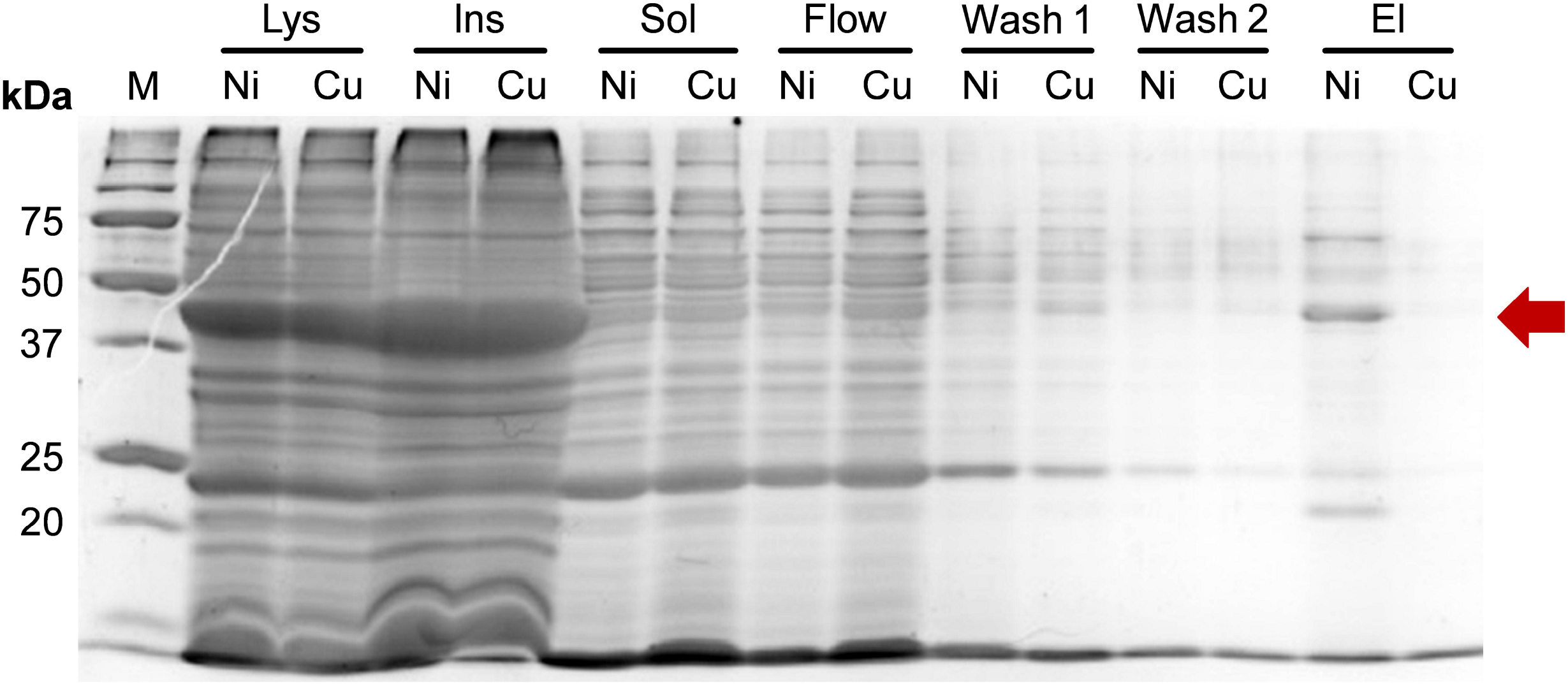
Binding of the fusion protein CusF-mCherry with IMAC. The protein was expressed in *E. coli* strains Rosetta 2 (DE3) and bind with Ni^2+^- or Cu^2+^-charged IMAC. M — Precision Plus Protein standards (Bio-Rad, Hercules, CA, USA). Lys — lysates after the expression, Ins — insoluble proteins, Sol — soluble protein fraction, Flow — flowthrow, Wash 1 — the 1^st^ wash, Wash 2 — the 2^nd^ wash, El — eluates. Ni and Cu — probes from Ni^2+^- or Cu^2+^-charged IMAC, respectively. Red arrow marks CusF-mCherry.

As observed for CusF–Efa PAP, the CusF–mCherry fusion protein was largely insoluble after expression and was nearly undetectable in the soluble protein fraction. However, the soluble fractions showed a faint red coloration, indicating the presence of small amounts of CusF–mCherry. We therefore proceeded to test the binding of the soluble material to IMAC resins charged with different metal ions. No CusF–mCherry was eluted from Cu^2+^-charged IMAC resin, as confirmed by SDS–PAGE, and the corresponding elution sample showed no visible red coloration. In contrast, a moderate band at the expected molecular weight of CusF–mCherry (approximately 40kD a) was detected after elution from Ni^2+^-charged IMAC resin, and this sample exhibited a clear red hue. Since the CusF–mCherry construct includes a C-terminal His-tag originating from the pET36b vector, this tag is likely responsible for the observed binding to Ni^2+^-charged IMAC. Thus, the CusF–mCherry fusion protein remained almost completely insoluble and did not bind to Cu^2+^-charged IMAC resin, while retaining the ability to bind Ni^2+^-charged IMAC via its C-terminal His-tag.

## 3. Discussion

Expression and purification of recombinant proteins are essential components of modern life sciences and biotechnology. High-yield production of soluble recombinant proteins facilitates purification as well as subsequent functional studies and practical applications. Although a broad range of strategies has been proposed for optimizing recombinant protein production, no universal solution has yet been found for consistently obtaining proteins in the desired soluble form. Protein solubility tags are no exception; they also possess limitations, such as large size (e.g., 42 kDa MBP or 55 kDa GST), thermal instability of mesophilic tags, interference with the activity of fused proteins, and challenges with tag removal due to incomplete proteolytic cleavage (e.g., by TEV protease). Among these limitations, unpredictable solubilization efficacy and variable expression yields remain particularly difficult to address, as the performance of a given tag cannot be reliably predicted for a new protein. In this context, parallel testing of multiple tags for a single protein is often the most effective strategy, underlining the need for empirical data on the performance of different tags.

We first employed CusF as a solubility tag for a putative poly(A) polymerase *from E. faecalis*. Its cognate proteins, *E. coli* PAP-1 and tRNA nucleotidyltransferase, are known to be aggregation-prone and unstable when stored in low-salt buffers [13,17,20]. Our previous attempts to solubilize Efa PAP using either MBP or GST fusions were unsuccessful, prompting us to use CusF, which had been reported as a promising solubility tag and was expected to facilitate purification via metal-chelate chromatography. However, the CusF–Efa PAP fusion protein remained almost completely insoluble, similar to unfused Efa PAP. The reason for this failure is unclear, as the molecular weight and pI of Efa PAP are comparable to those of GFP—the protein originally used in the proof-of-concept study demonstrating successful solubilization by CusF [10] (48.5 kDa and pI 5.29 vs 27 kDa and pI 5.9, respectively). Strain-specific effects are also unlikely, as we used the same BL21 (DE3) strain reported in earlier work, and the pLysS plasmid typically does not reduce recombinant protein solubility.

To further assess the solubilization potential of CusF, we used mCherry as a fusion partner. mCherry, a DsRed-derived fluorescent protein from *Discosoma* sp., shares the β-barrel fold of GFP despite having low sequence identity (22%). It is generally expressed in *E. coli* as a highly soluble protein, making it a suitable model. Surprisingly, the CusF–mCherry fusion was also nearly completely insoluble. Moreover, the chimeric protein did not bind to Cu^2+^-charged IMAC resin, contrary to previous reports. A plausible explanation is that CusF has relatively weak affinity for Cu(II) ions and preferentially binds Ag(I); for Cu(I), the K_d_ is approximately tenfold higher, and the reported K_d_ for Cu(II) is about fivefold lower than for Cu(I) [8]. The Cu^2+^ concentration on our IMAC resin may therefore have been insufficient for efficient CusF binding, unlike the conditions used by Cantu-Bustos et al. Regarding solubility, one might attribute poor solubility to insufficient ionic strength of the lysis buffer; however, mCherry is typically purified in 150–300 mM NaCl, and we used 300 mM NaCl in our experiments.

Several limitations of our study should be noted. First, we evaluated only two proteins as fusion partners for CusF, and both—Efa PAP and mCherry—are moderately sized proteins with acidic pI values (5–6). Other proteins, particularly basic proteins, might exhibit improved solubility when fused to CusF. Second, we did not systematically optimize lysis conditions (e.g., pH, ionic strength, detergents), which can strongly influence solubility; higher ionic strength may improve the solubility of CusF fusions. Third, we did not extensively test CusF–mCherry binding to Cu^2+^-charged IMAC under varied conditions. Adjustments such as pH optimization or the use of chaotropic agents might yield improved results. Nonetheless, despite these limitations, we believe the presented findings are valuable. Proof-of-concept studies typically emphasize successful cases, whereas negative or unsuccessful outcomes—such as failed solubilization attempts—are underreported. Thus, we decided to publish our unsuccessful attempts to use CusF as a solubility tag in order to share limitations of this approach which can be helpful to other scientists in planning their experiments. It should be also noted that in the context of rapid advances in AI-driven protein design, lack of empirical negative data may hinder the development and training of computational models, as the models will have only positive results.

In summary, we provide insights into the performance of CusF as a solubility tag using two model proteins—a putative poly(A) polymerase from *E. faecalis* and the fluorescent protein mCherry. In both cases, fusion with CusF did not improve solubility, and CusF–mCherry did not bind to Cu^2+^-charged IMAC resin. These results demonstrate that, similar to MBP, GST, and other protein tags, CusF does not guarantee soluble expression of its fusion partner, highlighting the importance of testing multiple tags to achieve optimal protein yields.

## 4. Materials and Methods

### 4.1. Search and selection of E. faecalis PAP coding sequence

Amino acid sequence of *E. coli* PAP-1 was used as a reference for search of potential bacterial PAPs using Protein BLAST against proteins from *Bacilli*. Criteria for cloning were length 400-600 a.a., the presence of all PAP structural domains (head, neck, body and leg), conservative positions corresponding to D69, D71, E108 involved in catalysis in PAP I *E. coli* [21], characteristic signature of bacterial PAPs [LIV][LIV]G[R/K][R/K]Fx-[LIV]h[HQL][LIV] [22].

### 4.2. Cloning of E. faecalis polyA polymerase, mCherry, CusF and chimeric proteins

The coding sequence of *E. faecalis* PAP-1 (GenBank: EOE33521.1) was amplified using pcnB-Efe-F/pcnB-Efe-R primers (Table 1) with NdeI and NotI restriction sites, allowing the in-frame ligation into the pET23a vector (Novagen, Madison, WI, USA). PCR was carried out using genomic DNA of *E. faecalis* as a template. The resultant 1.3-kbp DNA fragment and pET23a vector were digested with NdeI and NotI (SibEnzyme, Novosibirsk, Russia), ligated, and transformed into *E. coli* XL1-Blue cells according to the standard protocols [23]. The fidelity of the resulting recombinant plasmid named pPAP-Efa was confirmed by sequence analysis using primers pET-F and pET-R (Table 1) using the Big Dye Terminator kit 3.1 (Applied Biosystems, Waltham, MA, USA) and ABI 3730 genetic analyzer (Applied Biosystems, Waltham, MA, USA) in the laboratory of antimicrobial drugs at ICBFM SB RAS according to the manufacturer’s protocol. The coding sequences of mCherry and CusF were amplified (mCherry-F/mCherry-R and CusF-F/CusF-R (Table 1), respectively) and cloned using the same procedure into pET23a (NdeI and NotI) and pET36b (NdeI and KpnI), respectively, resulting in plasmids pmCherry and pCusF.

**Table 1.**
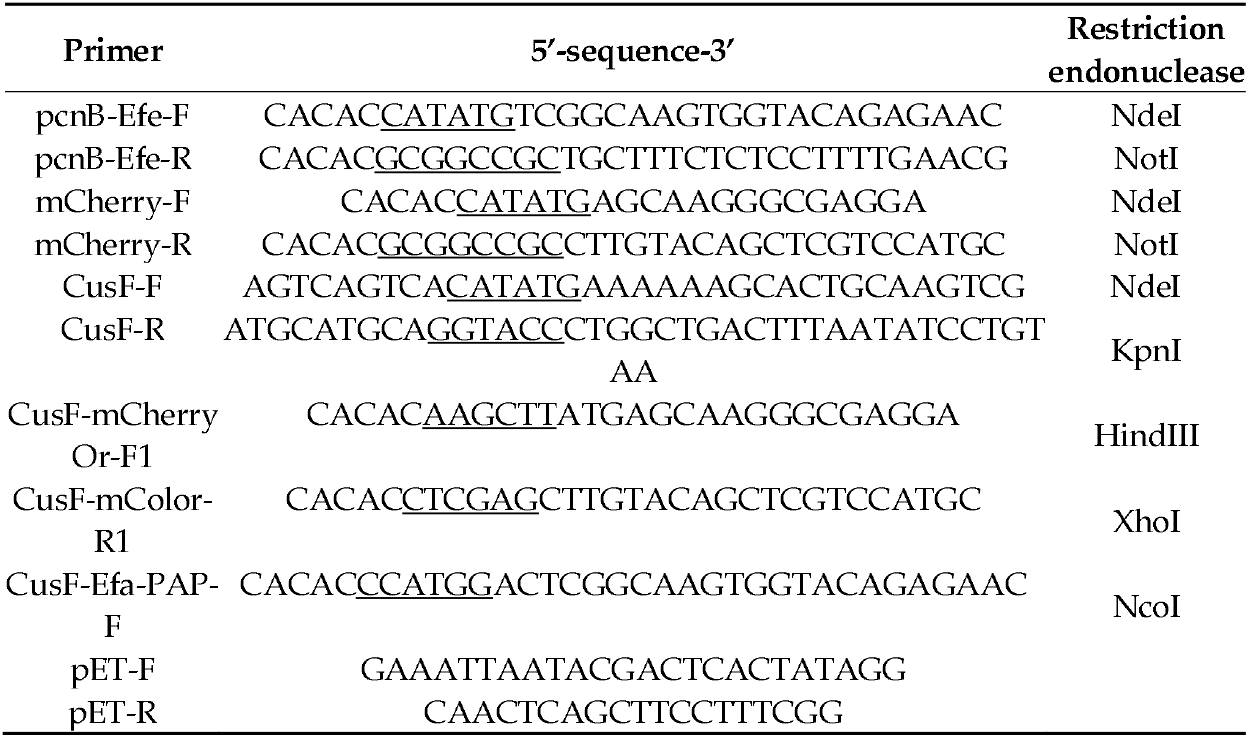
Oligonucleotide primers and probes.

For construction of the fusion proteins CusF-mCherry and CusF-Efa-PAP, the coding sequences of mCherry and Efa PAP were amplified (CusF-mCherryOr-F1/CusF-mCherryOr-R1 and CusF-Efa-PAP-F/mCherry-R (Table 1), respectively) and cloned into pCusF HindIII/XhoI and NcoI/NotI, respectively, resulting in plasmids pCusF-mCherry and pCusF-PAP-Efa.

### 4.3. Expression of a putative Efa PAP, CusF-Efa PAP, CusF-mCherry

A starter culture of *E. coli* BL21 (DE3) pLysS and (Promega, WI, Madison, USA) Rosetta 2 (DE3) (Novagen, WI, Madison, USA) strains harboring the plasmids pPAP-Efa, pCusF-mCherry, pCusF-PAP-Efa were grown to OD600 = 0.8 in LB medium with 100 μg/mL ampicillin (pPAP-Efa) or 50 μM kanamycin (pCusF-mCherry, pCusF-PAP-Efa) at 37 °C. In 4 1L flasks, 1 L of LB with 100 μg/mL ampicillin or 50 μM kanamycin was inoculated with 4 ml of the starter culture, and the cells were grown to OD600 = 0.6 at 37 °C. The expression of recombinant proteins was induced by adding IPTG up to 1 mM concentration. After induction for 12 h at 37 °C, the cells were harvested by centrifugation at 4,000 × g and stored at *−*70 °C.

### 4.4. Solubility test

To evaluate solubility of the fusion proteins in different *E. coli* strains, cell pellets after expression in 1 mL night cultures were lysed in 200 μl of a lysis buffer (50 mM Tris-HCl pH 8.0, 300 mM NaCl, 1 mM PMSF, 5% glycerol, 0.5% Triton X-100, 5 mM imidazole, 5 mM β-Mercaptoethanol). The cell pellets were resuspended in the lysis buffer following addition of lysozyme to a 1.0 mg/ml and incubation for 1 hour at 37 °C. The incubated probes were sonicated and centrifuged at 20,000 × g for 15 min. Soluble fractions were transferred into new tubes, and insoluble pellets were resuspended in 200 μl of the lysis buffer. Both soluble and insoluble fractions were analyzed using SDS-PAGE.

### 4.5. Binding of CusF-mCherry with Ni^2+^- and Cu^2+^-charged IMAC resin

For protein purification, the cell pellet was resuspended in 10 mL of the lysis buffer (50 mM Tris-HCl pH 8.0, 300 mM NaCl, 1 mM PMSF, 5% glycerol, 0.5% Triton X-100, 5 mM imidazole, 5 mM β-Mercaptoethanol) supplied by 1 mg/ml lysozyme, incubated for 30 min on ice followed by sonication. After lysis, the soluble fraction was separated by two consequent centrifugation steps at 20,000×g for 30 min following binding with Ni^2+^- or Cu^2+^-charged IMAC resin (Bio-Rad, CA, Hercules, USA) pre-equilibrated with the lysis buffer. The resins were washed twice by the lysis buffer following elution of bound proteins were eluted by an elution buffer (20 mM Tris-HCl pH 8.0, 300 mM NaCl, 10% glycerol, 0.5% Triton X-100, 0.5 M imidazole, 5 mM β-Mercaptoethanol). All the fractions from each step were analyzed by SDS-PAGE.

## 5. Conclusions

To sum up, CusF did not improve the solubility of either a putative *E. faecalis* poly(A)-polymerase or mCherry, and CusF-mCherry did not bind Cu^2+^-charged IMAC resin under the used conditions. These results demonstrate that CusF is not a universally effective solubility tag as other previously suggested to the same purpose proteins like MBP, GST, thioredoxin, etc. The presented results underline the importance of empirical testing multiple tags as being essential for achieving optimal recombinant protein yield.

## Author Contributions

Conceptualization, M.L.F. and I.P.O.; methodology, I.P.O., M.S.K. and M.A.S.; validation, I.P.O.; formal analysis, I.P.O. and M.A.S.; investigation, M.S.K. and M.A.S.; resources, M.L.F.; data curation, I.P.O.; writing—original draft preparation, I.P.O.; writing—review and editing, M.L.F.; visualization, M.S.K. and M.A.S.; supervision, I.P.O.; project administration, M.L.F.; funding acquisition, I.P.O. All authors have read and agreed to the published version of the manuscript.

## Institutional Review Board Statement

Not applicable. Informed Consent Statement: Not applicable.

## Data Availability Statement

Dataset available on request due to the restrictions (e.g., privacy, legal or ethical reasons).

## Abbreviations

Efa: Enterococcus faecalis
PAP: poly(A)-polymerase

## Disclaimer/Publisher’s Note

The statements, opinions and data contained in all publications are solely those of the individual author(s) and contributor(s) and not of MDPI and/or the editor(s). MDPI and/or the editor(s) disclaim responsibility for any injury to people or property resulting from any ideas, methods, instructions or products referred to in the content.

